# Identification of Tumor Antigens and Immune Subtypes of Acute Myeloid Leukemia for mRNA Vaccine Development

**DOI:** 10.1101/2022.10.25.513524

**Authors:** Fan Wang

**Affiliations:** Department of Hematology, Tongji Hospital, Tongji Medical College, Huazhong University of Science and Technology, Wuhan, Hubei, The People’s Republic of China

**Keywords:** mRNA vaccine, acute myeloid leukemia, tumor immune microenvironment, tumor antigen, Cancer vaccination

## Abstract

**Background:** Acute myeloid leukemia (AML) is a highly aggressive hematological malignancy, and there has not been any significant improvement in therapy of AML over the past several decades. The mRNA vaccines have become a promising strategy against multiple cancers, however, its application on AML remains undefined. In this study, we aimed to identify novel antigens for developing mRNA vaccines against AML and explore the immune landscape of AML to select appropriate patients for vaccination.

**Methods:** The genomic data and clinical information were retrieved from TCGA and GEO, respectively. The gene mutation data of AML were obtained from cBioPortal. GEPIA2 was used to analyze differentially expressed genes. The single cell RNA-seq database Tumor Immune Single-cell Hub (TISCH) was used to explore the association between the potential tumor antigens and the infiltrating immune cells in the bone marrow. Consensus clustering analysis was applied to identify distinct immune subtypes. The correlation between the abundance of antigen presenting cells and the expression level of antigens was evaluated using Spearman correlation analysis. The characteristics of the tumor immune microenvironment in each subtype were investigated based on single-sample gene set enrichment analysis. Weighted gene co-expression network analysis (WGCNA) was performed to identify co-expression modules and screen the hub genes.

**Results:** Five potential tumor antigens were identified for mRNA vaccine from the pool of overexpressed and mutated genes, including CDH23, LRP1, MEFV, MYOF and SLC9A9, which were associated with infiltration of antigen-presenting immune cells (APCs). AML patients were stratified into two immune subtypes Cluster1 (C1) and Cluster2 (C2), which were characterized by distinct molecular and clinical features. Patients with C1 subtype showed higher expression level of the five candidate genes and worse prognosis compared with C2 subtype. Moreover, C1 subtype demonstrated an immune-hot and immunosuppressive phenotype, while the C1 subtype had an immune-cold phenotype. Furthermore, the two immune subtype showed remarkably different expression of immune checkpoints, immunogenic cell death modulators and human leukocyte antigens. Finally, WCGNA identified a module and five hub genes that can be used for predicting the response of AML patients to mRNA vaccine.

**Conclusion:** CDH23, LRP1, MEFV, MYOF and SLC9A9 were potential antigens for developing AML mRNA vaccine, and AML patients in immune subtype 1 were suitable for vaccination.

## Introduction

Acute myeloid leukemia (AML) is a highly heterogeneous hematological malignancy arose from wild proliferation of undifferentiated myeloid blasts^1^, and it is the second most common type of leukemia^2^, occurring most in adults, with the average diagnosis age of 68^2^. The standard therapy regime of AML is consisted of induction phase and consolidation phase with chemotherapy and thereafter with possible allogeneic hematopoietic stem cell transplantation (HSCT) ^3^. However, this traditional therapy mode is commonly closely related to high toxicity and high risk of relapse, and there has not been any significant improvement in the treatment filed of AML over the past several decades^3^. Although the five-year survival rate for AML patients younger than 20 years old is around 60%-75%^4^, it is only dismal 3%-8% for patients over 60^5,6^. The truncated long-term survival probability of AML patients is mainly due to the common relapse after treatment^3^. Thus, novel effective treatment methods that can eradicate the residual AML cells is expected to be necessary for the cure of AML.

Immunotherapy, which can eliminate cancer cells without harming normal cells by establishing the immunosurvielling activity of the immune system against the cancer cells, could be a better therapeutic option for AML^7^. Immune checkpoint blockade (ICB) and chimeric antigen receptor T (CAR-T) cell therapy have gained tremendous success for the treatment of lymphoma but not for AML^8^. Cancer vaccination is a kind of immunotherapy that can induce specific T-cell responses potentially capable of specifically destroying cancer cells through introduction of certain cancer antigens^9^. The potential of these vaccine-induced cancer antigen-specific T cell responses to persist and establish immunological memory make it possible for the cancer vaccines to create a long-term protection against cancer recurrence^9^. Currently, there are several types of cancer vaccines explored in clinical trials, including DNA, peptides, dendritic cells (DCs) and RNA^10^. But for AML, only peptide and DC-based vaccines had been investigated to enhance leukemia-specific immune responses^11^. A phase II trial of WT1 (Wilms’ Tumor 1) peptide vaccination administered to 22 AML patients with a median age of 64 years after complete remission (CR1) showed that WT1 vaccinations in AML patients were safe and well tolerated, the median disease-free survival (DFS) from CR1 was 16.9 months, while the overall survival (OS) from diagnosis had not yet been reached but is poised to be ≥67.6 months^12^, which were superior to published data for similar patients treated with conventional postremission therapies or HSCT^13-15^. Another clinical trial of personalized vaccination of 17 postremission AML patients with a hybridoma of AML cells and autologous dendritic cells (DCs) demonstrated that the vaccination was well tolerated and 12 of 17 vaccinated patients (71%) remain alive without recurrence at a median follow-up of 57 months^16^. In 2021, the stunning success of the SARS-CoV-2 RNA vaccines for prevention of COVID-19 has led to considerable enthusiasm for mRNA vaccine^17^. There were a number of advantages for mRNA vaccine when compared with other treatments available in the clinic. Firstly, mRNA vaccine is quite safe since mRNA has no risk of insertional mutagenesis by gene integration which often happens to DNA vectors and mRNA can be easily degraded by normal cellular processes, the half-life of mRNA can be regulated using various RNA sequence modifications or delivery systems^18,19^. Secondly, mRNA vaccines can be manufactured *in vitro* in a rapid and inexpensive way without the need of complex process to produce antibody or viral vector drugs, hence it will save a lot of valuable time for patients with rapid-growing cancers^20,21^. Thirdly, *in vitro* mRNA production is highly versatile and efficient since it is very easy to modify the mRNA sequence for different target proteins^22^. Fourthly, unlike DNA vector or protein drugs, there is almost no anti-vector immunity for mRNA^23^. Thus, mRNA vaccine is highly practicable for targeting tumor-specific antigens as a promising immunotherapy scheme. Currently, dozens of phase 1/2 clinical trials are underway to prove the effectiveness of mRNA vaccines in patients with various types of cancer, including melanoma, lung cancer, pancreatic cancer, colorectal cancer, ovarian cancer, prostate cancer and relapsed/refractory lymphoma^24^. However, no effective mRNA vaccine for AML has been developed so far.

In this study, we aimed to identify novel antigens for developing mRNA vaccines against AML and explore the immune landscape of AML to choose appropriate patients for vaccination. Five neoantigen candidates associated poor prognosis and positively correlated to the infiltration of antigen-presenting cells (APCs) were identified in AML patients. Based on the clustering of immune-related genes, two robust immune subtypes were defined and then validated in independent cohorts. The two immune subtypes were associated with distinct molecular, cellular, and clinical features. Finally, two functional modules closely correlated to immune subtypes were screened through gene co-expression network analysis (WGCNA). These findings might provide a theoretical basis for developing mRNA vaccine against AML and facilitate choosing appropriate patients for vaccination.

## Materials and Methods Data acquisition

The normalized RNA sequencing (RNA-seq, TPM) data of 70 normal bone marrow samples from the GTEx (Genotype-Tissue Expression) database and 173 AML patients from the TCGA (The Cancer Genome Atlas) database with the corresponding clinical data were extracted from the “TCGA TARGET GTEX” dataset of Xena (https://xenabrowser.net/datapages/). The RNA-seq data with clinical information of 146 AML patients were downloaded from the GEO database (https://www.ncbi.nlm.nih.gov/geo/, GSE147515) and were used as an independent cohort for external validation. The GSE147515 dataset is consisted of transcriptomics data of 1523 samples from 11 datasets covering 10 AML cytogenetic subgroups, which were then merged with the transcriptomic data of 198 healthy bone marrow samples. The gene mutation data of AML were acquired from cBioPortal (https://www.cbioportal.org/) based on the AML samples in TCGA, the mutant genes in AML were screened with the R package “maftools” and the corresponding chromosome position of genes were plotted with the R package “RCircos”.

### GEPIA Analysis

Differentially expressed genes (DEGs) of AML was analyzed using the online database Gene Expression Profiling Interactive Analysis (GEPIA, http://gepia2.cancer-pku.cn/) based on samples from the TCGA and the GTEx databases. The differential analysis was performed by comparing AML to paired normal samples with the “limma” package, and the chromosomal distribution of over- or under-expressed genes were plotted in differential expressed genes with a cutoff of |Log2FC| > 1 and q-value < 0.01.

### ESTIMATE Analysis

The immune and stromal scores of each AML sample were computed by using the R package “estimate”. The AML samples were then divided into high and low score groups according to the median value of their stromal and immune scores, the immune-related DEGs and stromal-related DEGs between the high score and low score groups were screened by the “limma” package, and they were then intersected by the “venn” package.

### Prognosis Analysis

To evaluate the prognostic value of potential AML antigens, overall survival (OS) and disease-free survival (DFS) analysis was performed using the Kaplan-Meier method with a 50% (Median) cutoff for the gene expression. Log-rank P-values < 0.05 were considered statistically significant.

### TISCH analysis

The single cell RNA-seq database Tumor Immune Single-cell Hub (TISCH, http://tisch.comp-genomics.org/) was used to investigate the distribution of potential tumor antigens in the infiltrating immune cells of the bone marrow. The TISCH database GSE116256 was divided into 22 cell clusters and 13 major cell types, the individual gene expression was visualized on various immune cell types.

### Identification and validation of the immune subtypes

RNA sequencing data of 173 AML patients from the TCGA cohort were transformed using log2(x+0.001), consensus clustering analysis was applied to identify distinct immune subtypes based on 2,483 immune-related genes by using the “ConsensusClusterPlus” package of R. Cluster sets varied from 2 to 9 and the optimal k value as the number of clusters was defined by assessing the consensus matrix and the consensus cumulative distribution function. The overall survival between the immune subtypes were compared by using Kaplan–Meier analysis. In addition, the correlation between immune subtypes and clinical features including gender, cytogenetic risk (favorable, intermediate, poor), age, survival status, white blood cell, blasts of bow marrow or peripheral blood and status of mutated genes were explored. To evaluate the reliability of the identified immune subtypes, another independent AML cohort from the GEO database (GSE147515) was used as the validation group with the same algorithm.

### Differential analysis of HLA, ICPs, TMB, and ICDs

The distribution of tumor mutational burden (TMB) and tumor stemness index mRNAsi were compared between the immune subtypes. Moreover, the expression level of human leukocyte antigen (HLA) genes, immune checkpoint genes and immunogenic cell death modulators (ICDs) were compared among different immune subtypes.

### Immune cell infiltration analysis

The immune cell infiltration of TCGA samples based on gene expression profiling were calculated by the current acknowledged algorithms including XCELL, QUANTISEQ, Microenvironment Cell Populations-counter (MCPcounter), EPIC, CIBERSORT-ABS and CIBERSORT as already published^25^. The correlation between the abundance of APCs and the expression level of the potential tumor antigens was evaluated using Spearman correlation analysis. The difference of immune cell infiltration between the AML immune subtypes were analyzed using t-test. Immune cells were filtered with *P* < 0.05 and visualized utilizing the R package “pheatmap”. Moreover, a total of 28 immune gene sets^26^ representing diverse adaptive and innate immunity were quantified for their enrichment scores within the respective AML samples using single-sample gene set enrichment analysis (ssGSEA) by the R package “GSVA”. The ssGSEA score were then compared between the AML immune subtypes and the results were plotted by the “ggplot2” package.

### Weighted Gene Co-expression Network Analysis

The R package “WGCNA” was used to identify co-expression modules by clustering the samples. Gene modules were then examined by dynamic hybrid cut. Univariate and multivariate Cox regression models were used for analyzing the independent prognostic value of the gene modules. Gene Ontology (GO) and Kyoto Encyclopedia of Genes and Genomes (KEGG) analysis were performed to annotate the functions of the identified modules with the “clusterProfiler” package. A protein-protein interaction (PPI) network was built using the STRING website (https://string-db.org/) with the cutoff of 0.7 to assess the relationship between the eigengenes of the finally identified gene module, and the results were visualized using the Cytoscape software (https://cytoscape.org/, version 4.0.4), the hub genes were screened using the “cytoHub” plug-in.

## Results

### Identification of potential antigens for mRNA vaccine of AML

To identify potential antigens of AML, we first screened out 6595 significantly overexpressed genes (Log2 fold change >1) that could encode tumor-associated antigens (Fig. 1A). Considering tumor-associated antigens are significantly related with gene mutations, a total of 406 mutated genes with mutation number ≥ 3 in AML were filtered, and their corresponding positions in human chromosomes were labeled as shown in Fig. 1B. The mutation landscape of the top 30 genes with the highest mutation frequency were shown (Fig. 1C, Figure S1). Since immune infiltration is an important determinant of the effect of cancer immunotherapy, 997 genes were obtained by intersecting 1290 immune related DEGs and 1189 stromal related DEGs (Fig. 1D). Finally, 9 genes including CACNA2D3, CDH23, SLC9A9, MYOF, TNFSF10, CXCL16, CYBB, MEFV, LRP1 were screened out for encoding potential tumor-associated antigens in AML through the intersection of overexpressed genes, mutant genes, and immune infiltration related DEGs, the numbers and intersections of which were shown in Fig. 1E. The related genes were listed in Supplementary Table1.

**Fig. 1.**
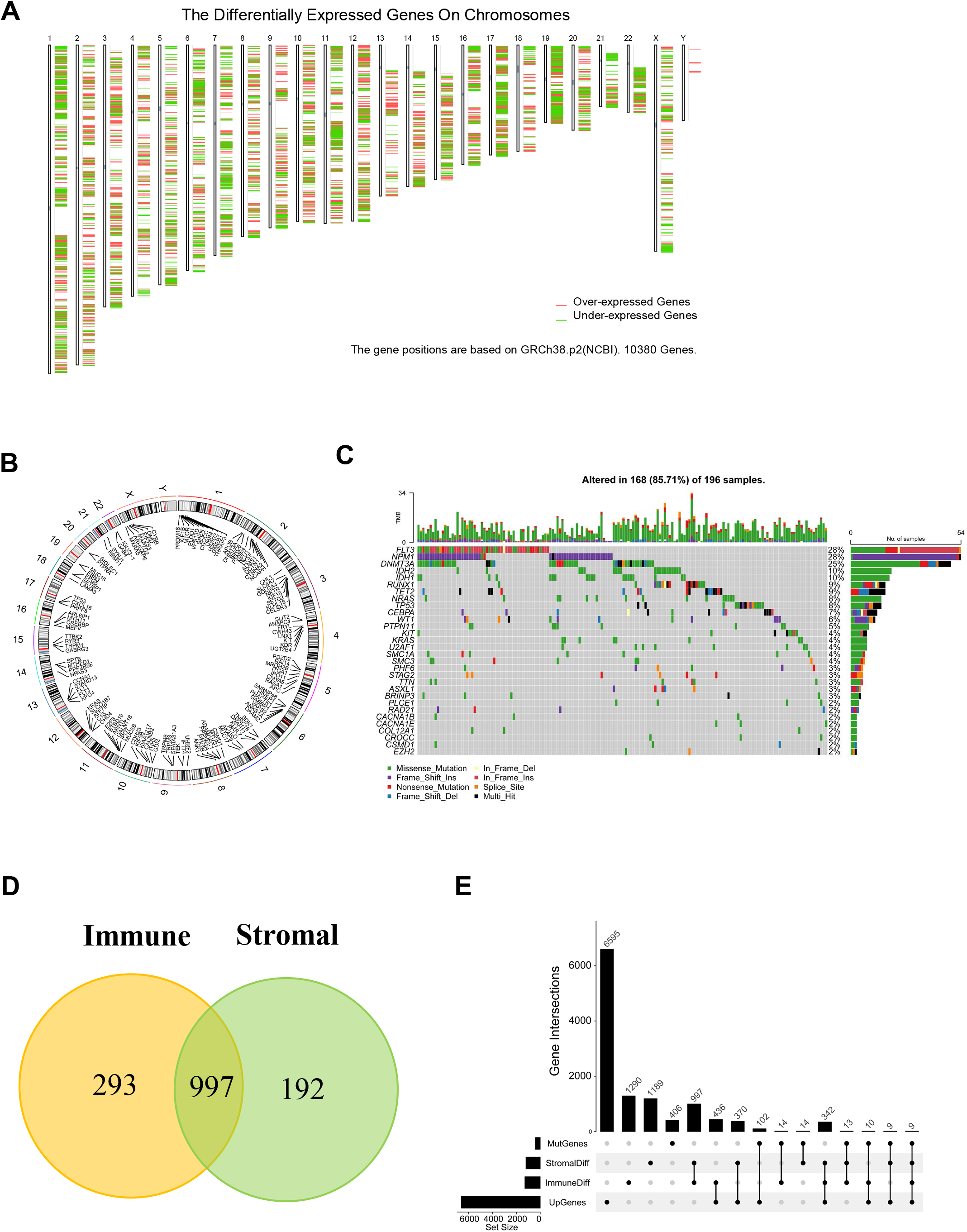
Identification of potential tumor antigens for mRNA vaccine in AML. (A) The chromosomal distribution of upregulated and downregulated genes in AML. (B) A circle plot of chromosomal distribution of mutated genes in AML. (C) The waterfall plot of the distribution of the top 30 mutated genes in AML. (D) A venn diagram of immune related DEGs and stromal related DEGs. (E) The upset plot of the intersections of genes screened under different conditions. AML, acute myeloid leukemia; MutGenes, mutated genes; StromalDiff, differentially expressed genes among low and high stromal score groups; ImmuneDiff, differentially expressed genes among low and high immune score groups; DEGs, differently expressed genes; UpGenes, upregulated genes in AML.

### Evaluation of the prognosis value and correlation with APCs of the nine potential tumor antigens in AML

To evaluate the prognosis value of the 9 potential tumor antigens, survival analysis was performed to further filtered prognostically associated antigens as candidates for mRNA vaccine development in AML. A total of 5 genes (CDH23, LRP1, MEFV, MYOF, SLC9A9) significantly correlated with the DFS of AML were identified, of which two (CDH23, MYOF) were significantly related to the OS (Fig. 2). Patients with elevated expression of CDH23, LRP1, MEFV, MYOF and SLC9A9 showed significant shorter DFS compared to the lower expression group (Fig. 2 F-J). The five antigens also demonstrated inferior OS in the high expression group, however, only CDH23 and MYOF showed significance (Fig. 2 A-E). Thus, 5 candidate genes were identified that are critical for the progression of AML. Considering professional APCs including dendritic cells (DCs), macrophages and B cells playing significant roles in the onset of protective immunity and effectiveness of mRNA vaccines by capturing and cross-presenting the antigens to activate T cells, the immune cell infiltrations in AML samples were estimated through XCELL, QUANTISEQ, MCPCOUNTER, EPIC, CIBERSORT-ABS and CIBERSORT algorithms^25^, respectively. Spearman correlation analysis showed that the expression level of CDH23, LRP1, MEFV, MYOF and SLC9A9 were significantly positively associated with infiltrations of myeloid dendritic cell, naïve B cell, macrophage, macrophage M1 and macrophage M2, except for some negative correlation with naïve B cell using the CIBERSORT algorithm (Fig. 3). These findings suggest that the five identified tumor antigens can be processed and presented by APCs to trigger a robust immune response.

**Fig. 2.**
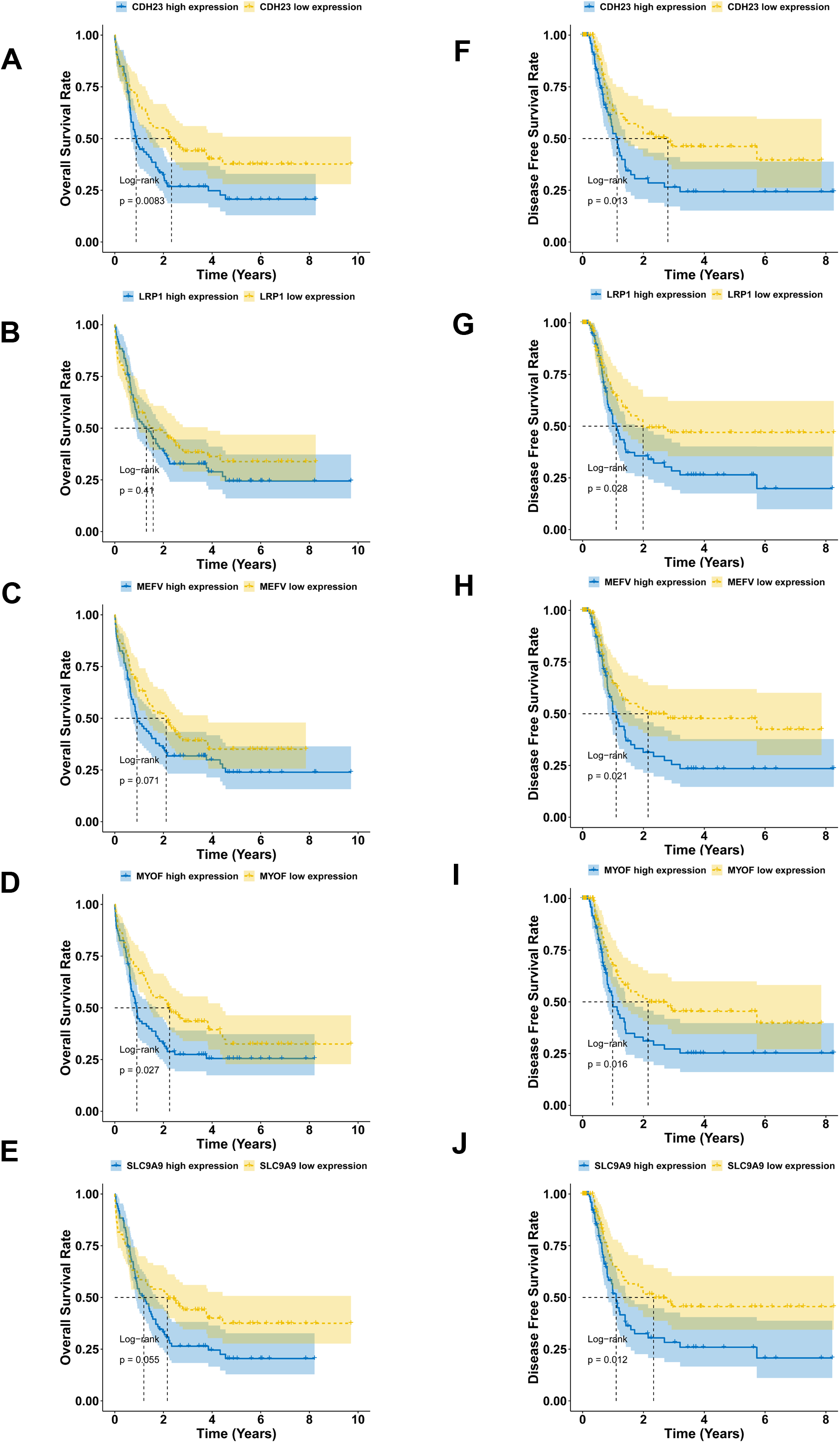
The prognostic value of the nine potential antigens for mRNA vaccine in AML. Kaplan-Meier curves showed the overall survival (A-E) and disease-free survival (F-J) of AML patients in the different expression levels of (A,F) CDH23, (B,G) LRP1, (C,H) MEFV, (D,I) MYOF and (E,J) SLC9A9.

**Fig. 3.**
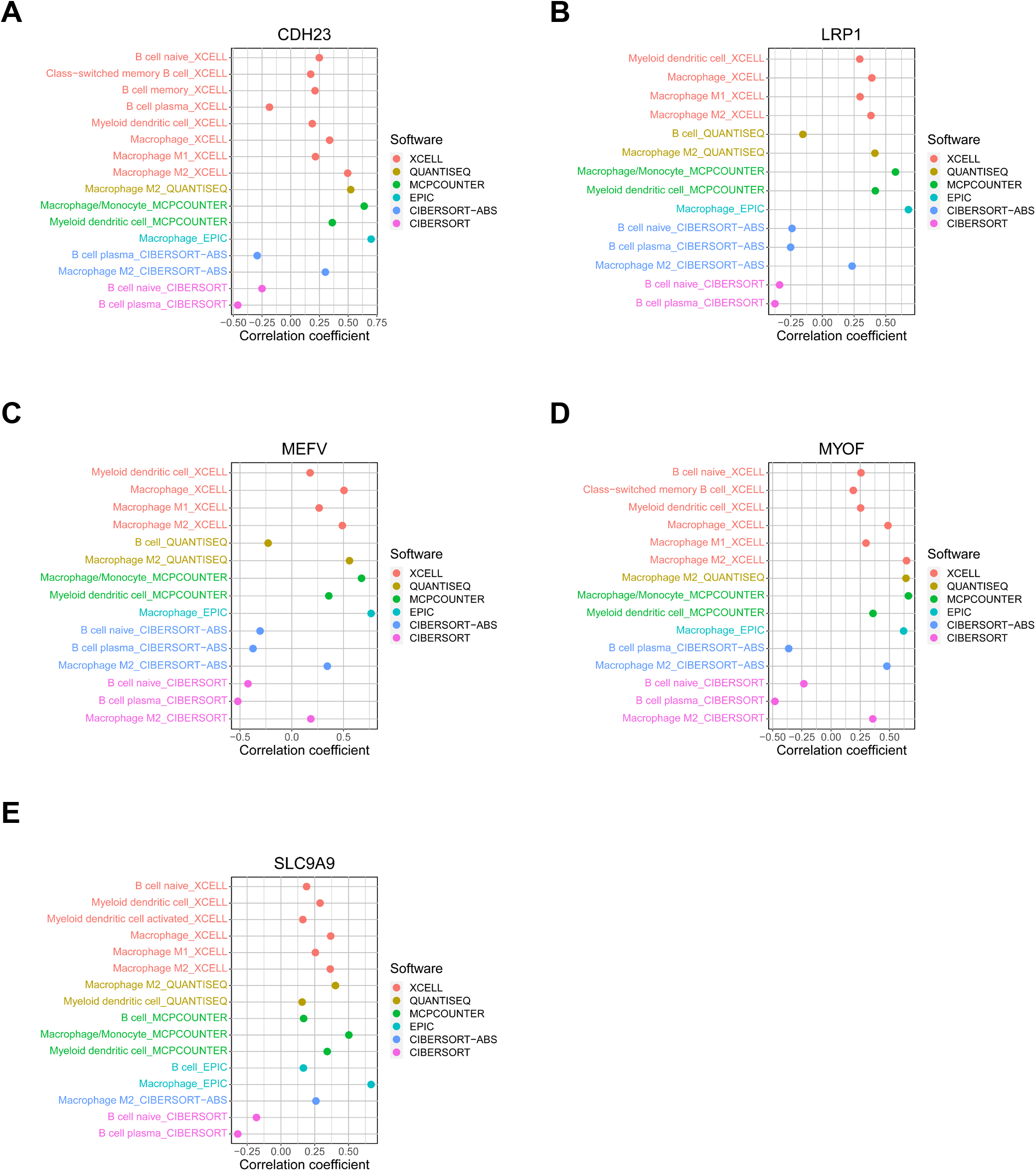
Correlations between the five candidate genes and immune cell infiltrations of TCGA AML samples. The infiltrating immune cells of TCGA AML samples were estimated by using current acknowledged methods such as XCELL, QUANTISEQ, MCPcounter, EPIC, CIBERSORT-ABS and CIBERSORT. Spearman correlation analysis was performed to evaluate the correlation between the five candidate genes (A) CDH23, (B) LRP1, (C) MEFV, (D) MYOF, (E) SLC9A9 and the immune cells. Only data with *P* < 0.05 were shown.

Furthermore, single cell analysis of various immune cell types of the bone marrow from AML patients in the TISCH dataset GSE116256 demonstrated that the five candidate genes were highly expressed in macrophages (Fig. 4). Taken together, CDH23, LRP1, MEFV, MYOF and SLC9A9 were identified as potential tumor-specific antigens for mRNA vaccine in AML.

**Fig. 4.**
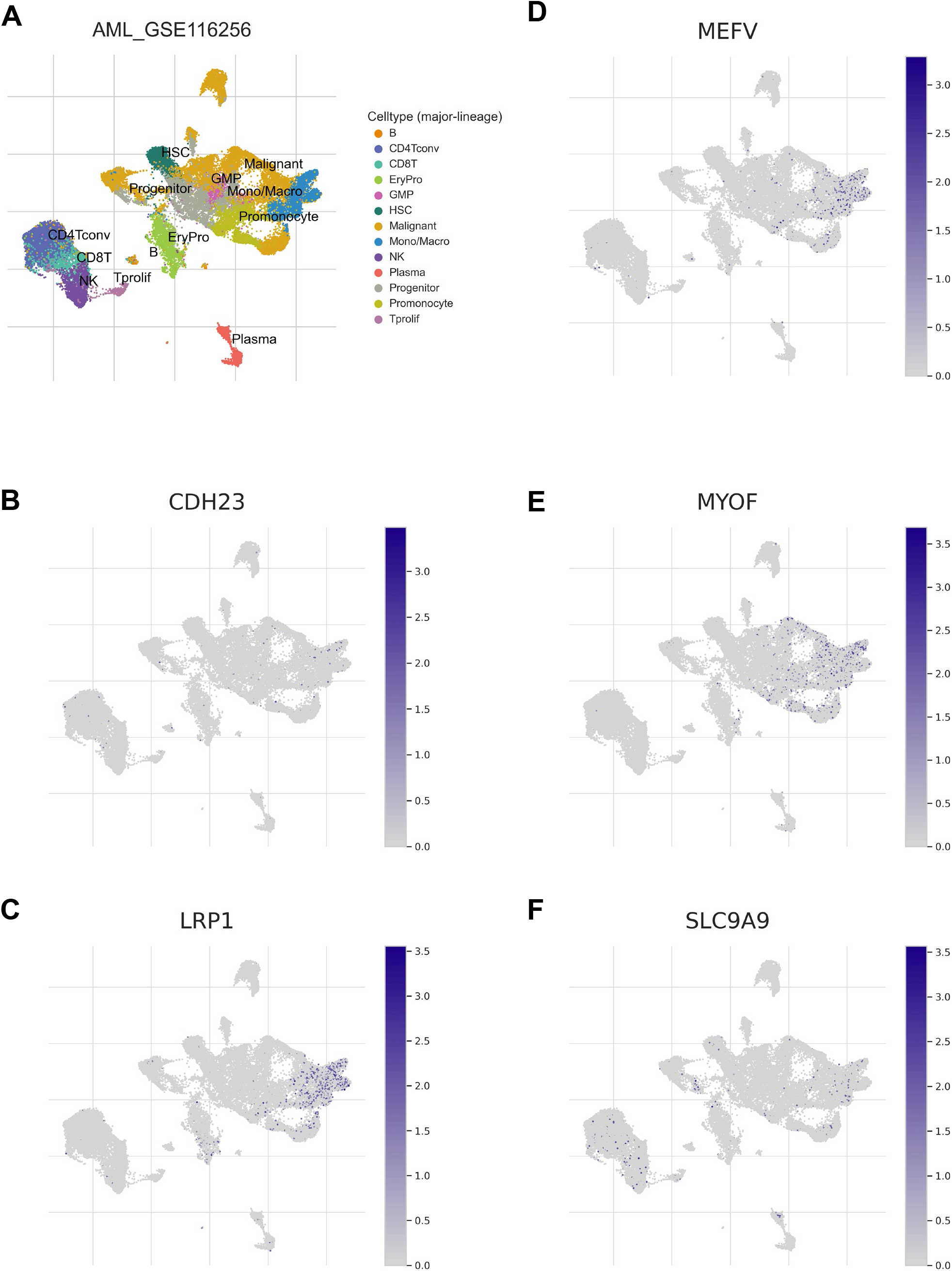
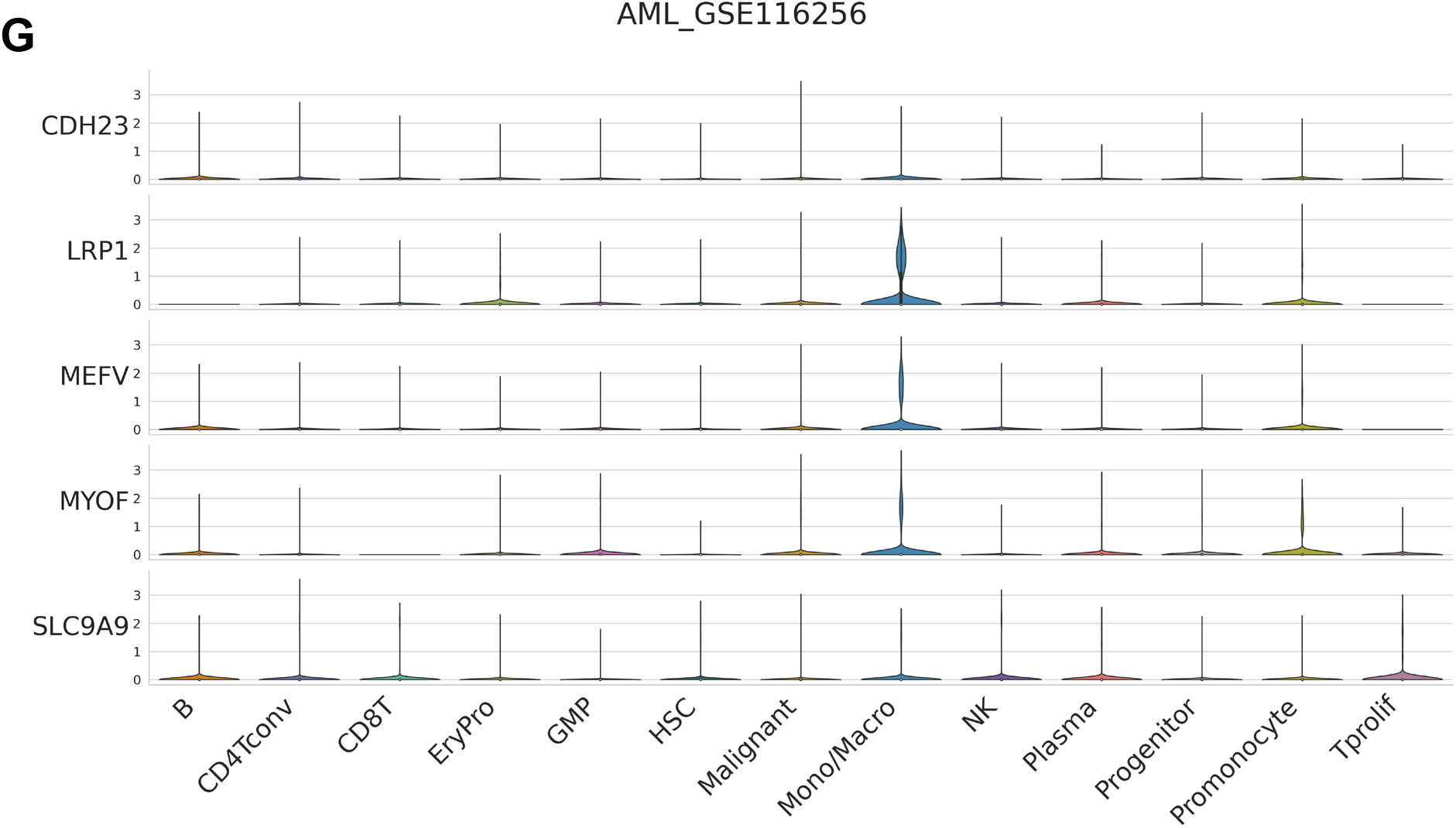
Analysis of the expression level of the five candidate genes in various cell types in GSE116256 from the TISCH database. **(A)** Annotation of the major cell types in the GSE116256 dataset. **(D-F)** Expression levels of (B) CDH23, (C) LRP1, (D) MEFV, (E) MYOF, (F) SLC9A9 in a variety of cell types of the GSE116256 dataset. **(G)** Comparison the expression level of CDH23, LRP1, MEFV, MYOF and SLC9A9 in different cell types of the GSE116256 dataset.

### Identification of immune subtypes of AML

Since the tumor microenvironment (TME) of AML is heterogeneous, it is important to identify patients suitable for vaccination with the mRNA vaccine. A consensus clustering analysis was performed in the TCGA AML cohorts based on the 2483 immune gene profiles. According to the corresponding cumulative distribution function (CDF) and function delta area of k value (Fig. 5A, B), the subtype clustering appeared to be stable while k =2, thus two robust immune subtypes (Cluster 1, Cluster 2) were obtained (Fig. 5C). Survival analysis demonstrated a significant difference between the two different immune subtypes, where Cluster1 (C1) group displayed a significantly worse prognosis than the Cluster2 (C2) group (*P* < 0.05, Fig. 5D). Consistent with the data obtained from the TCGA cohort, C1 group showed inferior prognosis compared to the C2 group in the GEO cohort as well (*P* < 0.001, Fig. 5E), suggesting the stability and reproducibility of the results. Therefore, immunotyping can be employed to predict prognosis of AML patients, and patients in the C2 group will have better prognosis. Furthermore, the gene expression profile of the five potential tumor antigens and the clinicopathological characteristics were compared among the two immune subtypes. The clinical characteristics including gender, cytogenetic risk (favorable, intermediate, poor), age (<60 or >=60 years), WBC (white blood cell) (<100 * 10^9^/L, >= 100 * 10^9^/L), blasts of BM (bow marrow) or PB (peripheral blood) (<50%, >= 50%), status of mutated genes (RUNX1, NRAS, KRAS, TET2, DNMT3A, TP53, IDH1, NPM1, WT1, FLT3) and cluster group were plotted in a heatmap. It was found that in the C1 group, there are significantly more cases with the clinical features of older age (>=60 years), less blasts of BM or PB, poor cytogenetic risk, more mutations of RUNX1 and TP53 (Fig. 5F). There are more samples with higher expression level of CDH23, LRP1, MEFV, MYOF and SLC9A9 found in the C1 subtype, indicating patients in this subtype may have higher specificity for mRNA vaccine treatment in AML. For the GEO cohort, the mutation data were lacking, but the expression level of the candidate genes were also found to be higher in the C1 group (Fig. 5G).

**Fig. 5.**
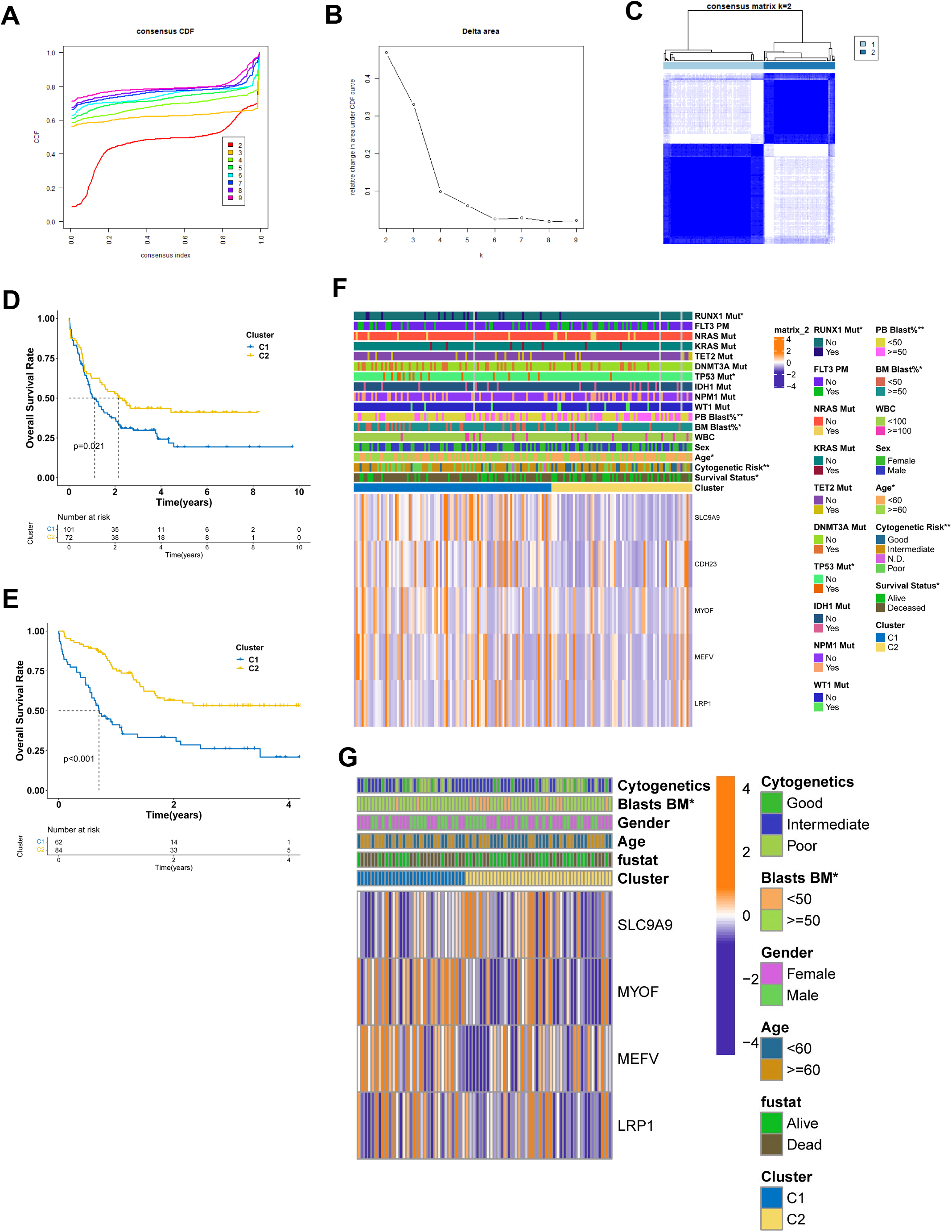
AML clustering based on the 2483 immune genes. (A) Cumulative distribution function (CDF) curve and (B) delta area plot of the immune-related genes in the TCGA cohort (k = 2∼9). (C) Consensus clustering matrix for k = 2 in the TCGA cohort. (D) Kaplan–Meier overall survival curves of the two clusters in TCGA cohort. (E) Kaplan–Meier overall survival curves of the two clusters in GEO cohort. (F-G) Differential analysis of clinicopathological characteristics and expression level of the five potential vaccine antigens in the two subgroups of the TCGA cohort (F) and GEO cohort (G). C1, Cluster1; C2, Cluster2; BM, bow marrow; PB, peripheral blood; WBC, white blood cell.

### Correlation analysis of immune subtypes and tumor mutational burden

Mutations in cancer cells will produce new epitopes of self-antigens (neoantigens), which can elicit antitumor immunity mediated by cytotoxic T cells. Cancer cells with high tumor mutational burden (TMB) are tended to possess more candidate neoantigens for vaccine development ^27^. At first, the general mutation analysis of TCGA-AML was shown in Figure S1. Then the TMB and mutations were compared between the two immune subtypes, but no significant differences were found (Figure S2A & B). Moreover, 20 genes including FLT3, NPM1, RUNX1 and TP53 were most frequently mutated in both subtype (Figure S2C). Cancer stem cell characteristics are related to the development of AML, but no significant variation was observed between the stemness of the two subtypes by quantifying the cancer stemness of each tumor samples using the stem cell-associated index mRNAsi (Figure S2D, *P* = 0.051).

### Correlation of immune subtypes with and immune modulators and HLA

Given the important roles of immune checkpoints (ICPs) and immunogenic cell death (ICD) modulators in cancer immunity, which can affect the efficacy of mRNA vaccine, the expression levels of ICPs and ICD were assessed in the two immune subtypes. A total of 59 ICPs-related genes were detected in the TCGA and GEO cohorts. Twenty-eight genes were distinctly expressed in the two subtypes in the TCGA cohort, and C1 had significantly higher expression of CD40, CD226, ICOS, CD80, BTN3A1, ICOSSLG, KIR2DL4, ADORA2A, BTN2A2, CD28, BTN2A1, CD274, CEACAM1, CD40LG, TNFRSF9, TNFSF14, CTLA4, KIR2DL3, TIGIT, KIR2DL1, KIR2DS4, PDCD1 (PD-1), CD160, BTLA, KIR3DL1, PDCD1LG2 (PD-L2) and CD27 (Fig. 6A). However, 25 genes were differentially expressed in the two subtypes in the GEO cohort, and C1 had higher C10orf54, CD86, LGALS9, SIRPA, TNFSF4 and TNFSF9 expressions (Fig. 6B). Moreover, 33 ICD-related genes were detected in both TCGA and GEO cohorts, of which 16 genes and 11 genes were differentially expressed in the two immune subtypes of the TCGA and GEO cohort, respectively (Fig. 6C & D). C1 had significant upregulation of PRF1, CD8B, EIF2AK3, FOXP3, IL1R1, CASP1, NT5E, CD4, CXCR3, LY96, IL10, IFNG and CD8A in the TCGA cohort (Fig. 6C), and higher expression of BAX, CASP1, IFNGR1, IL17RA, MYD88, NLRP3, P2RX7 and TLR4 in the GEO cohort (Fig. 6D). Human leukocyte antigens (HLA) play critical roles in antigen processing and presentation, the expression level of 54 HLA genes were assessed in the two immune subtypes. The expression levels of 24 HLA genes were significantly elevated in C1 compared to C2, C1 showed higher expression of HLA-DOB, HLA-DQB2, HLA-F, HLA-DQA1, HLA-DMB, PSMB8, HLA-DMA, HLA-B, HLA-C, HLA-E, HLA-DRB6, HLA-DPB1, TAP2, HLA-DPA1, HLA-K, HLA-L, PSMB9, HLA-DRA, HLA-DOA, TAP1, MICD, HLA-DRB1, MICE and HLA-H in the TCGA cohort (Fig. 6E). Collectively, the response of patients in C1 group to the mRNA vaccine treatment could be more effective and promising.

**Fig. 6.**
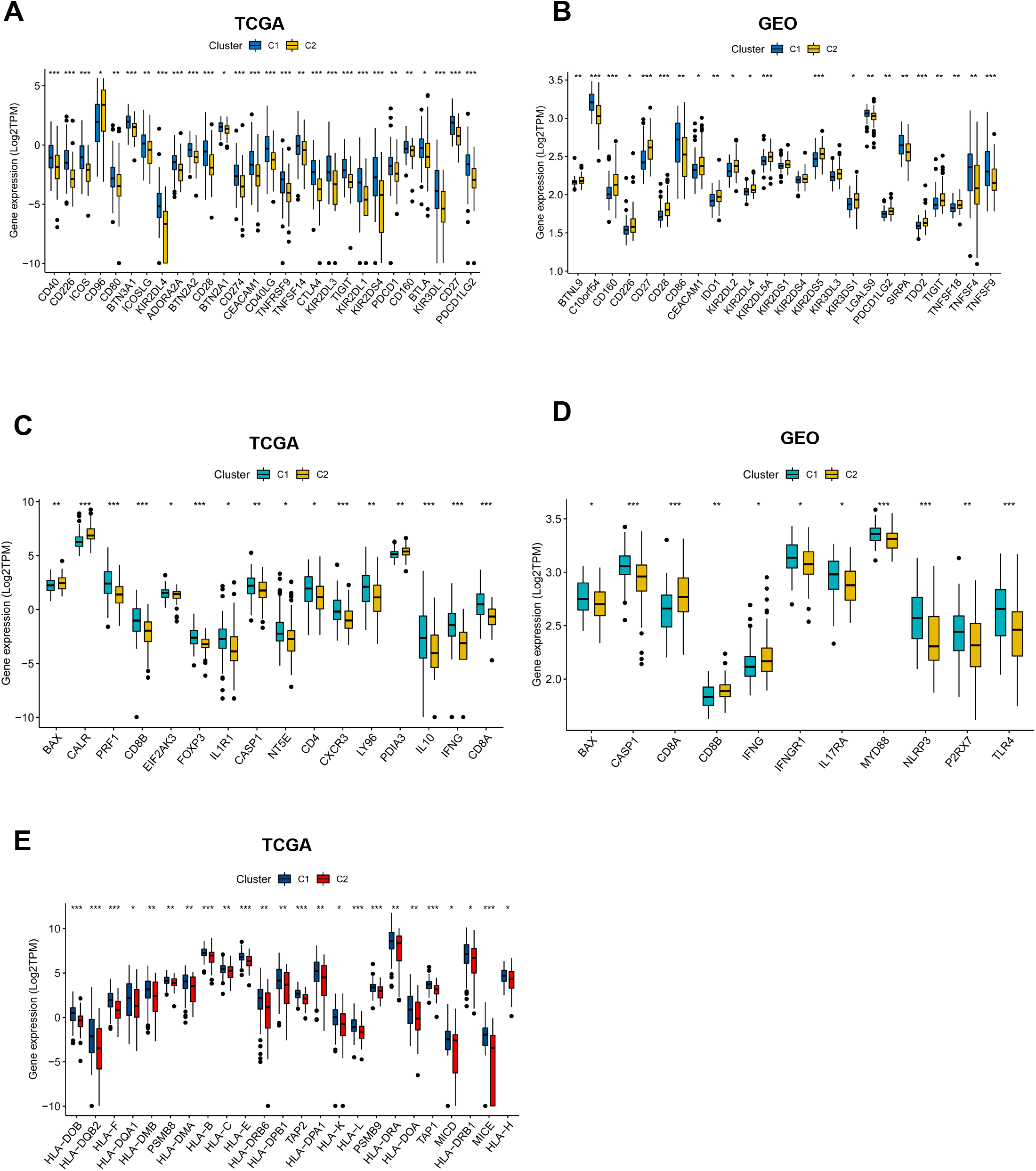
Association of immune subtypes with ICPs, ICD modulators and HLA genes in AML. (A, B) Differential expression of ICPs genes between the two immune subtypes in (A) TCGA and (B) GEO cohorts. (C, D) Differential expression of ICD modulator genes between the two immune subtypes in (C) TCGA and (D) GEO cohorts. (E) Difference in the expression of HLA genes between the two immune subtypes in TCGA cohort. HLA, human leukocyte antigens; ICPs, immune checkpoints; ICD, immunogenic cell death. * *P* < 0.05, ** *P* < 0.01, *** *P* < 0.001, and **** *P* < 0.0001.

### Difference of immune microenvironment features in immune subtypes

The tumor immune microenvironment (TME) is extraordinarily important for the effectiveness of mRNA vaccine, thus the immune status between the two subtypes were analyzed. The immune cell infiltrations in TCGA AML samples were estimated through XCELL, QUANTISEQ, MCPCOUNTER, EPIC, CIBERSORT-ABS and CIBERSORT algorithms, as previously published^25^. The results showed that C1 had significantly higher infiltration of CD4+ T cell, CD8+ T cell, myeloid dendritic cell, naïve B cell, monocyte, macrophage, macrophage M2, NK cell and regulatory T cell (Fig. 7A). C1 also demonstrated higher immune score than C2 by using the XCELL algorithm (Fig. 7A). The immune cell abundance in the two immune subtypes were further defined by determining the scores of 28 previously reported immune signature gene sets in both TCGA and GEO cohorts using ssGSEA^28^. The immune cell components were found to be remarkably distinct between the two subtypes (Fig. 8B & C). C1 had significantly higher scores in activated B cell, activated CD4+ T cell, activated CD8+ T cell, central memory CD4+ T cell, effector memory CD8+ T cell, gamma delta T cell, immature B cell, T follicular helper cell, type 1 T helper cell, type 2 T helper cell, activated dendritic cell, CD56dim natural killer cell, myeloid-derived suppressor cells (MDSC), natural killer cell, natural killer T cell and plasmacytoid dendritic cell in the TCGA cohort (Fig. 7B). For the GEO cohort, compared to C2, the C1 also showed higher level of central memory CD4 T cell, central memory CD8 T cell, eosinophil, immature dendritic cell, macrophage, mast cell, monocyte and plasmacytoid dendritic cell (Fig. 7C). Thus, C1 is an immune-hot and immunosuppressive phenotype, while C2 is an immune-cold phenotype, and C1 could be more promising to respond to the mRNA vaccine. The immune landscape based on the two immune subtypes can be used identify suitable patients for personized mRNA vaccine therapy.

**Fig. 7.**
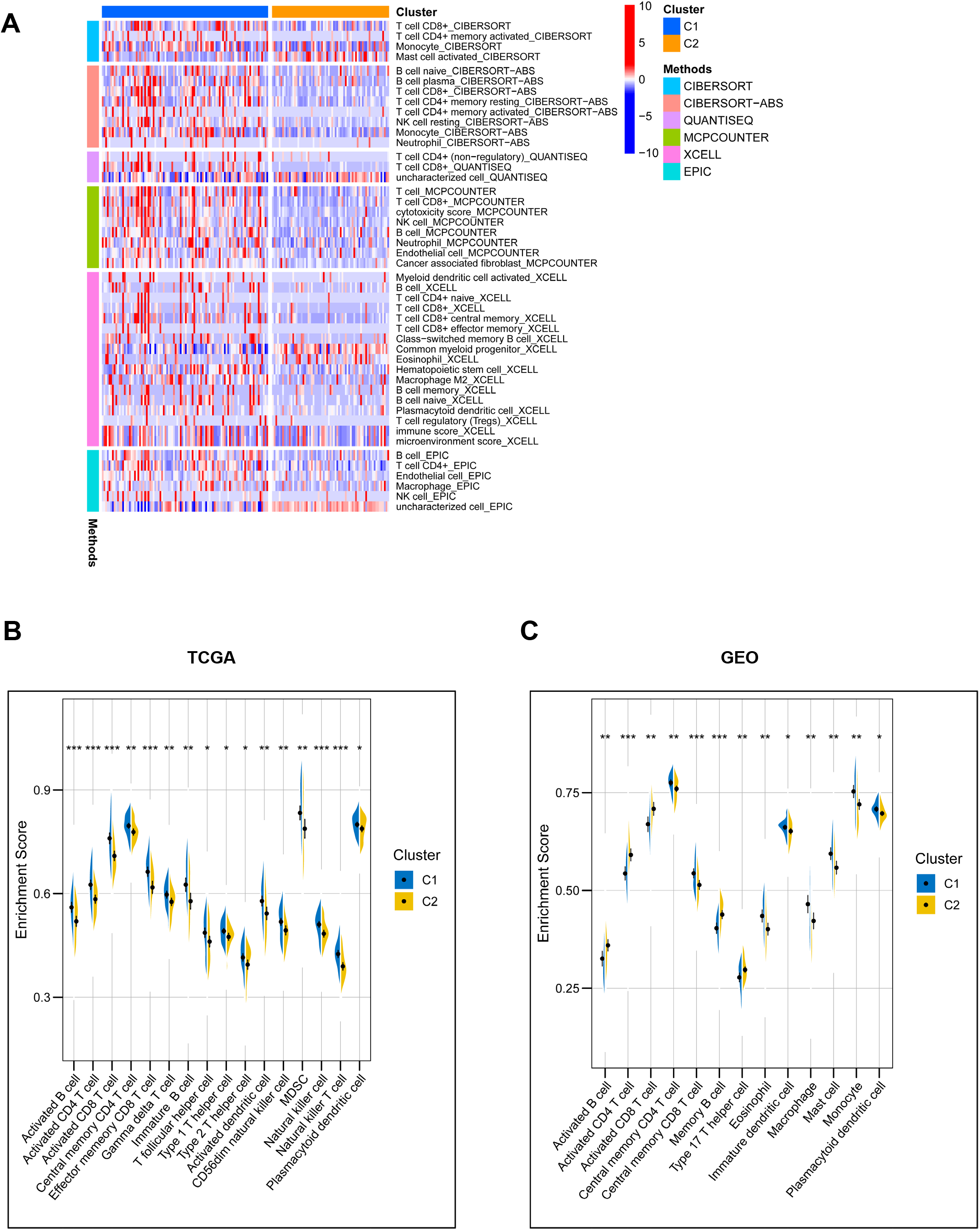
Difference of immune microenvironment characteristics in two immune subtypes. (A) The immune cell infiltrations in TCGA AML samples were estimated by using XCELL, QUANTISEQ, MCPCOUNTER, EPIC, CIBERSORT-ABS and CIBERSORT algorithms, the infiltration level were compared between the two groups C1 and C2. Only data with *P* < 0.05 were shown in the heatmap. (B, C) The difference in immune scores based on the results of single-sample gene-set enrichment analysis (ssGSEA) of 28 immune-related signatures between two immune subtypes in TCGA cohort (B) or GEO cohort (C). * *P* < 0.05, ** *P* < 0.01, *** *P* < 0.001, and **** *P* < 0.0001.

**Fig. 8.**
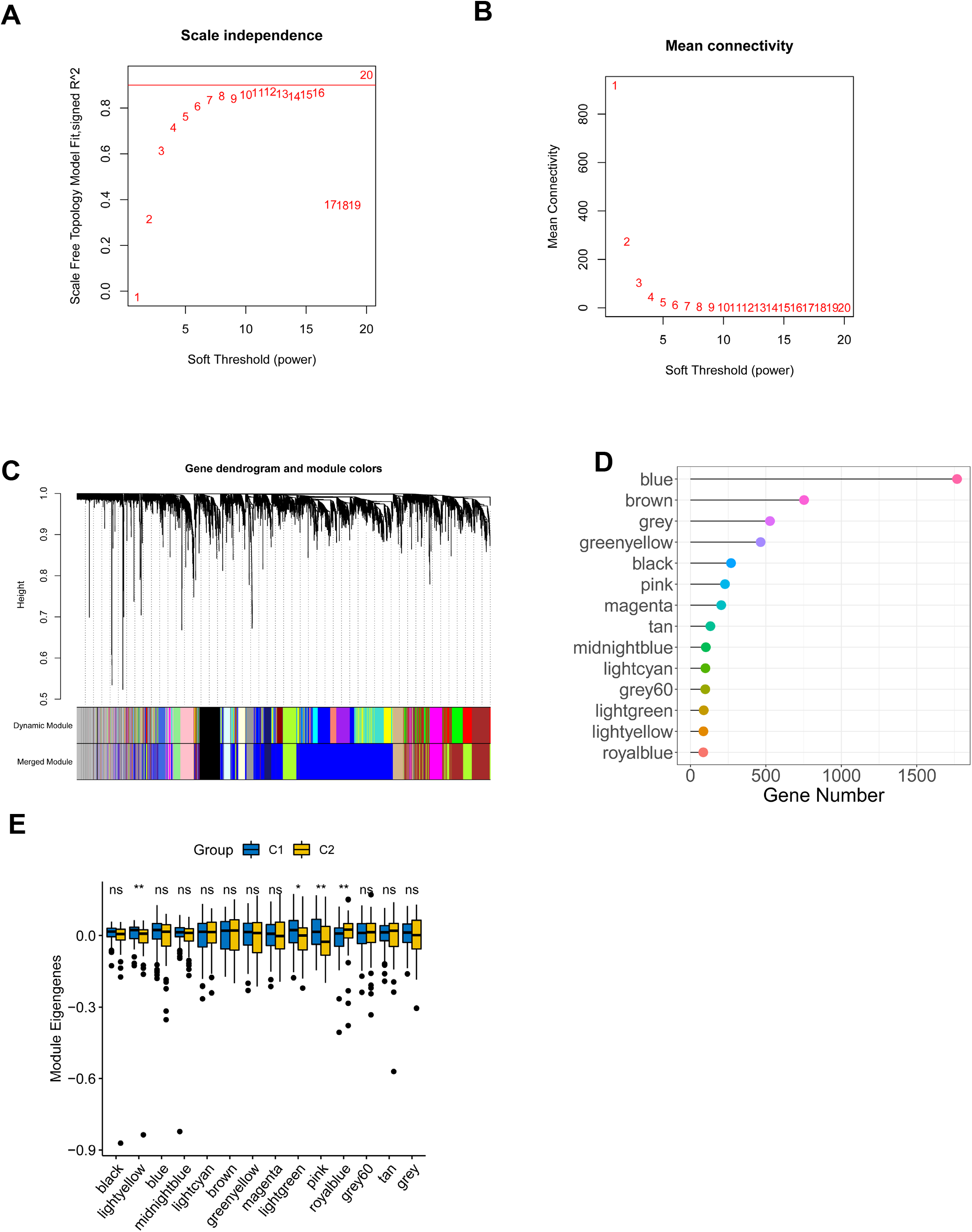
Identification of co-expression modules of TCGA AML cohort by WGCNA. (A) Scale-free fit index for various soft-thresholding powers (β). (B) Mean connectivity for various soft-thresholding powers. (C) Differentially expressed genes were clustered using hierarchical clustering with a dynamic tree cut and merged based on a dissimilarity measure (1-TOM). (D) Gene numbers of each module. (E) Differential distribution of each module in two AML immune subtypes. * *P* < 0.05; ** *P* < 0.01; ns, not significant.

### Identification of co-expression modules and hub genes

WGCNA was used to identify co-expression modules by clustering the samples with a soft threshold of 8 for scale-free network (Fig. 8A and B). The representation matrix was then converted to adjacency and next to a topological matrix. The average-linkage hierarchy clustering procedure was applied with a minimum of 30 genes for each network in line with the standard of a hybrid dynamic shear tree. Eigengenes of each module were determined and the close modules were consolidated into a new one (height = 0.25, deep split = 4 and min module size = 60) (Fig. 8C). Consequently, 14 co-expression modules with 4904 transcripts were acquired (Fig. 8D). The eigengenes of the 14 modules were then analyzed in the two immune subtypes, C1 showed significantly higher eigengenes in black, lightgreen and pink modules than C2 (Fig. 8E). Moreover, the univariate Cox regression analysis revealed that the expression of genes in the brown, grey, magenta, pink and tan modules were significantly associated with the poor prognosis of AML patients (Fig. 9A). Multivariate analysis further indicated that only brown and pink modules were independent survival prognostic factors (Fig. 9B). Furthermore, Gene ontology (GO) enrichment analysis and Kyoto Encyclopaedia of Genes and Genomes (KEGG) pathway analysis suggested that ribonucleoprotein complex biogenesis and ribosome were significantly enriched in brown module (Fig. 9 C & E), while pink model was significantly enriched with MHC protein complex binding, positive regulation of leukocyte mediated immunity, macrophage activation, B cell proliferation, positive regulation of T cell differentiation, antigen processing and presentation signaling (Fig. 9 D & F). Thus, only pink module is related with immune genes. Consistently, patients with higher scores of genes clustered into brown (Fig. 9G) and pink (Fig. 9H) modules had poor survival compared to those with lower scores in the TCGA cohort. Therefore, mRNA vaccine could be effective in patients with the highly expressing genes clustered into the pink module. Finally, 5 hub genes including CD4, ITGB2, ITGAM, FCGR2A and TLR2 were identified in the pink module (Fig. 9I), which can be potential biomarkers for predicting the response of AML patients to mRNA vaccine.

**Fig. 9.**
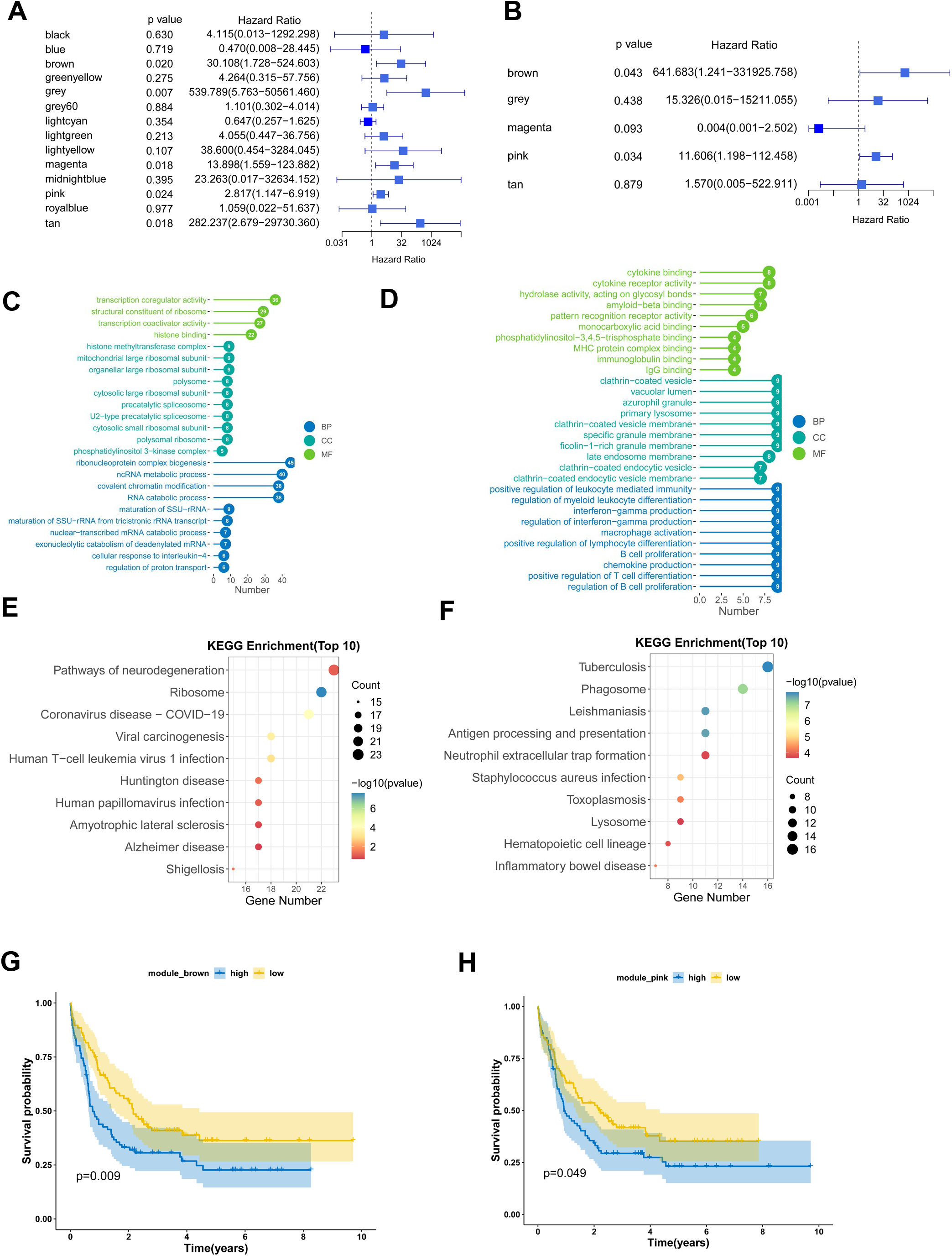
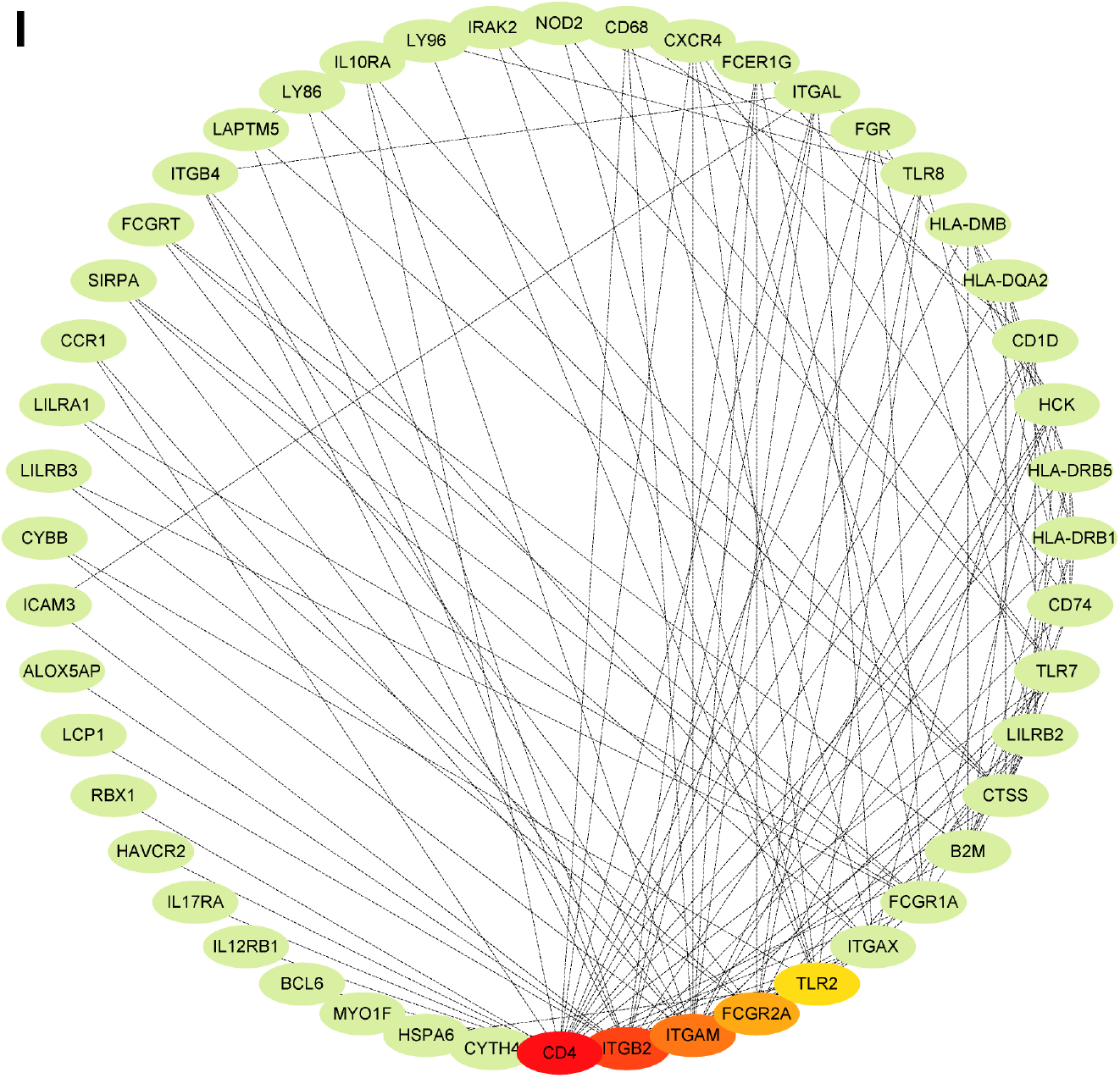
Identification of immune hub genes of AML. (A) Forest plot of univariate analysis of 14 identified modules of AML. (B) Forest plot of multivariate analysis of the 5 prognosis-related modules of AML. (C) Bar plot for GO enrichment of the brown module. (D) Bar plot for GO enrichment of the pink module. (BP: biological process, CC: cellular component, MF: molecular function). (E) Dot plot for top 10 KEGG enrichment of the brown module. (F) Dot plot for top 10 KEGG enrichment of the pink module. The dot size and color intensity represent the gene count and enrichment level respectively. (G) Differential prognosis in brown module with high and low scores stratified by the mean. (H) Differential prognosis in pink module with high and low scores. (I) Hub Genes of the pink Module, which is shown with the Degree Sorted Circle layout of Cytoscape, with the nodes’ color reflective of the level of connectivity within the network.

## Discussion

AML is a highly aggressive cancer that are mostly treated with chemotherapeutic drugs^1^. But this traditional therapy method is commonly closely related to high toxicity and high risk of relapse, and there has not been any significant improvement in the overall survival of AML over the past several decades^3^. Especially, old AML patients still have a very poor prognosis^4^. The mRNA vaccines have become trending in cancer immunotherapy since its successful application in preventing COVID-19^17^. However, there was rare studies exploring in treatment of AML with mRNA vaccine. Considering that the shortened overall survival of AML patients is by and large related to the relapse after chemotherapy^3^, the potential of the mRNA vaccine-boosted antigen-specific T cell responses to persist and establish post-treatment immunological memory will provide the chance of long-term protection against AML recurrence. In this study, the prospective tumor-associated antigens in AML were screened out through the intersection of aberrantly expressed genes, mutant genes and immune infiltration related DEGs, of which CDH23, LRP1, MEFV, MYOF and SLC9A9 were significantly correlated with the DFS of AML. Further analysis showed that those five antigens were significantly positively correlated with the infiltration of APCs, including dendritic cells, B cells and macrophages. And single cell analysis of various immune cell types of the bone marrow from AML patients demonstrated that the five candidates were highly expressed in macrophages. Thus, CDH23, LRP1, MEFV, MYOF and SLC9A9 were identified as potential tumor-specific antigens for mRNA vaccine development in AML. They can be processed and presented by APCs to cytotoxic T cells triggering a robust immune response that will attack the tumor cells. CDH23 (Cadherin-23) is a member of the calcium-dependent cell adhesion glycoproteins that constitutes the cadherin superfamily^29^. Studies showed that CDH23 played a critical role in cancer progression. For instance, CDH23 was upregulated in breast cancer and was associated with the early metastasis^30^, methylated depletion of CDH23 was related with poor prognosis in Diffuse large B-cell lymphoma (DLBCL) ^31^, germline mutations of CDH23 were linked with both familial and sporadic pituitary adenoma^32^. LRP1 (Low density lipoprotein receptor-related protein 1, also known as CD91) is a ubiquitously expressed endocytic receptor belonging to the low-density lipoprotein receptor (LDLR) superfamily, of which LRP1 is the most multifunctional one^33^. Studies have demonstrated that LRP1 is involved in two prime physiological activities: endocytosis and modulation of signaling pathways^34^, indicating that LRP1 may play multiple roles in tumorigenesis and tumor progression. It was reported that LRP1 inhibition induced suppression of Notch signaling and reduced tumorigenesis in leukemia models^35^. Another study reported that the expression of membrane-associated proteinase 3 (mP3) in AML blasts inhibits T cell proliferation via direct LRP1 and mP3 interaction, suggesting the importance of LRP1 in regulating the immunity environment in AML^36^. MEFV (Mediterranean fever) encoding pyrin is expressed in certain white blood cells including neutrophils, eosinophils and monocytes that is involved in regulation of inflammation and in fighting infection by interacting with the cytoskeleton^37^. MEFV mutations lead to reduced or malformed pyrin that cannot perform its presumed role in controlling inflammation, result in an inapplicable or extended inflammatory response^37^. In a study with a colitis mouse model, it was found that MEFV was required for inflammasome activation and IL18 maturation, which can avert colon inflammation and tumorigenesis^38^. Moreover, there is evidence that activation of autoinflammatory pathways including MEFV in the clonal cells of myelodysplastic syndrome and AML may be related with neutrophilic dermatoses^39^. MYOF (Myoferlin) is a member of the Ferlin family involved in membrane trafficking, membrane repair and exocytosis^40^. Accumulating evidence have revealed that MYOF is an oncogene that is overexpressed in a variety of cancers^41^. MYOF drives the progression of cancer by promoting tumorigenesis, proliferation, migration, epithelial to mesenchymal transition, invasiveness and angiogenesis^41^. For example, depletion of MYOF in breast cancer can significantly reduce tumor development and metastatic progression that was linked with degradation of the epidermal growth factor (EGF) receptor (EGFR) ^42^. Clinically, MYOF overexpression is associated with poor prognosis in patients with breast cancer, lung cancer, and pancreas cancer^43^. SLC9A9 (also called NHE9), a member of the Na+/H+ exchanger (NHE) superfamily, is a transmembrane protein that localizes mostly on the recycling endosome and plays a crucial role in regulating the pH of endosomes^44^. It has been reported that SLC9A9 was involved in attention autism and deficit hyperactivity disorder (ADHD) ^45^. Recently, some studies revealed that SLC9A9 gene was implicated in cancer as well. In esophageal squamous cell carcinoma, elevated expression level of SLC9A9 were associated with cancer advancement and inferior prognosis^46^. In glioblastoma, SLC9A9 was overexpressed and promoted the proliferation and invasiveness of glioblastoma cells through the activated EGFR signaling pathway^47^. In colorectal cancer, SLC9A9 was upregulated and can promote progression of colorectal cancer, which is closely related with EGFR pathway^48^. In addition, high level of SLC9A9 was involved in poor prognosis in colorectal cancer^48^. In this study, CDH23, LRP1, MEFV, MYOF and SLC9A9 were also markedly associated the prognosis of AML patients, which had not been reported previously. Moreover, to the best of our knowledge, this is the first study to screen antigens for developing mRNA vaccine for AML patients.

AML is a hematological malignancy with a high degree of genetic and immunological heterogeneity^49^, which has fundamental implications for the efficacy of immunotherapy and will restrict the widespread application of mRNA vaccine in AML patients. Therefore, it is essential to stratify AML patients according to their immune profiling which will be used to identify appropriate patients for mRNA vaccination. In this study, two reproducible immune subtypes of AML (C1 and C2) were identified based on the expression profile of 2483 immune genes. The two immune subtypes were associated with distinctive clinical features, for example, in the C1 immune subtype, there were significantly more patients with older age (>=60 years), less blasts of bone marrow or peripheral blood, poor cytogenetic risk, mutations of RUNX1 and TP53. Moreover, the C1 subtype showed higher expression level of the five candidate antigens compared to the C2 subtype. Furthermore, the C1 subtype displayed an inferior prognosis than C2 subtype in both TCGA and GEO cohorts. These data suggested completely different immunological and molecular patterns between the two immune subtypes.

Successful antitumor effect of the mRNA vaccine requires optimal tumor microenvironment (TME) ^50^. Within the TME, natural killer (NK) cells, macrophages and neutrophils of the innate immune system are required for immediate recognition and attacking of tumor cells, while APCs cross-present the antigens through interaction with T cell receptor (TCR) to activate T cells of the adaptive immune system, which are ultimately responsible for killing the cancer cells and eradicating the tumor^51^. A phase II trial of a multivalent WT1 peptide vaccine administered to 22 AML patients after first CR reported the immunologic responses were documented in 64% of patients, including increased CD4+ T cell proliferation and CD8+ T cell IFN-γ secretion, and the immunologic response seemed to be associated with improved survival outcomes^52^. In another phase II study, DCs electroporated with WT1 mRNA were administered to 30 patients with AML at very high risk of relapse, the long-term clinical response and outcome were linked with the induction of WT1-specific CD8+ T cell reaction^53^. In this study, C1 subtype had a significantly higher infiltration of antigen-presenting cells (APCs) and anti-cancer lymphocytes, such as CD4+ T cell and CD8+ T cell, indicating that the mRNA vaccine should trigger a more potent immune response against AML cells within the C1 subtype than the C2 subtype and AML patients with the C1 subtype should be more promising to respond to the mRNA vaccine. Thus, the immune landscape based on the two immune subtypes can be used identify suitable patients for personized mRNA vaccine therapy. However, C1 subtype was also more abundant with immunosuppressive immune cells such as regulatory T cells (Tregs) and myeloid-derived suppressor cells (MDSCs), two vital factors in tumor immune escape, which could compromise the effect of anti-cancer immune response mediated by the immune effector cells. Therefore, C1 subtype was considered as an immune-hot and immunosuppressive phenotype, whereas C2 was an immune-cold phenotype, which may also explain why the C1 subtype had a worse prognosis than C2. Evidence showed that ‘cold’ tumors are refractory to immunotherapy and ‘hot’ tumors are more responsive to immunotherapy^54^, suggesting that the mRNA vaccine could be more successful for AML patients with the C1 subtype. However, to ensure long-term protection against tumor relapse, theoretically, memory T cells induced via mRNA vaccination should persist for a long-time following tumor eradication. Thus, the memory T cell populations should be investigated in preclinical models and in AML patients in the future studies.

The mRNA vaccine transported via lipid nanoparticle enters the APCs to encode the target tumor antigens, which can be presented on the surface of APCs by MHC (also known as HLA in human) to evoke an antitumor response via interactions with TCR^24^. In this study, C1 subtype showed higher expression of HLA, which may have a greater response to the therapy with mRNA vaccine. It is now appreciated that the immunosuppressive TME substantially hinders the efficacy of mRNA vaccines^55^. The implementation of immune checkpoint inhibition (ICI) such as anti-CTLA-4, anti-PD1 and anti-PDL1 antibodies, successfully reprogramed one or more immunosuppressive signals in the TME to allow T cell to unleash its function, substantially increasing response rates and even leading to potential cures^56^. In the clinical trials, mRNA vaccines have been applied to treat solid tumors, including non-small cell lung cancers, melanoma, prostate cancer, and glioblastoma^9^. The combination of mRNA vaccines with ICI may further improve their antitumor efficacy. For example, a study of melanoma patients intranodally administered mRNA vaccine that encoded ten personalized neoantigens, showed an extraordinary vaccine-specific anticancer T cell response and a sustained progression-free survival. One relapsed patient exhibited complete response following anti-PD1 therapy. Another relapsed patient did not show to anti-PD1 therapy but turned out to have complete loss of HLA class I presentation on tumor cells due to β2M deficiency^57^. In another study of melanoma patients intravenously administered with mRNA vaccine consisting of four melanoma-associated antigens (NY-ESO-1, MAGE-A3, tyrosinase and transmembrane phosphatase with tensin homology (TPTE)), strong T cell responses were found to be correlated with durable clinical responses when combined with anti-PD1 therapy in patients with anti-PD1 resistance^58^. These results demonstrate that successfully induced anticancer responses require the co-delivery of ICI in addition to the mRNA vaccination since ICI can overcome immunological tolerance to tumor antigens. In the current study, the C1 subtype had significantly higher expression of ICPs such as CTLA4, ICOS, CD28, CD274, PDCD1 (PD-1) and PDCD1LG2 (PD-L2). Moreover, C1 subtype also had significant upregulation of immunogenic cell death (ICD) modulators including CD8B, EIF2AK3, FOXP3, IL1R1, CASP1, CD4, CXCR3 and IL10. These findings suggest that AML patients with the C1 subtype may get a better prognosis by administering with mRNA vaccination along with or in parallel with immune checkpoint inhibition. However, clinical benefit can only be confirmed in further research such as clinical trials.

Furthermore, the mRNA vaccination approaches may be combined with CAR-T cell that was pretty successful in treating hematological malignancies such as diffuse large B-cell lymphoma^59^. Combination of CAR-T cell therapy with cancer vaccines could increase the durability of CAR-T cells and even establish a lasting protection against cancer relapse. Currently, some researchers are investigating the effect of combinations of therapeutic CAR-T cell therapy with DC vaccines and RNA vaccines^60,61^. However, such endeavor for AML therapy is still lacking.

## Conclusions

In conclusion, CDH23, LRP1, MEFV, MYOF and SLC9A9 were potential antigens for AML mRNA vaccine development, and patients in immune subtype C1 were suitable candidates for such vaccination. This study will provide theoretical justification for constructing AML mRNA vaccine and selecting appropriate AML patients for vaccination.

## Data availability

The datasets analyzed during this study are available at TCGA, TARGET, UCSC Xena and GEO database (https://portal.gdc.cancer.gov/, https://ocg.cancer.gov/programs/target/, https://xenabrowser.net/, and https://www.ncbi.nlm.nih.gov/geo/; GSE147515).

## Author Contributions

F.W. came up with the conceptualization, methodology, software, data curation, writing, visualization, validation and funding acquisition.

## Declaration of Competing Interest

The author(s) declare that they have no conflict of interest.

## Compliance with Ethical Standards

This article does not contain any studies with human participants or animals performed by any of the authors.

## Funding

This work was supported by the National Natural Science Foundation of China, No. 82070174. The funder had no role in study design, collection, analysis and interpretation of data, decision to publish, or preparation of the manuscript.

## Acknowledgments

The results published here are in part based upon data generated by the Therapeutically Applicable Research to Generate Effective Treatments (https://ocg.cancer.gov/programs/target) initiative, phs000465. The data used for this analysis are available at https://portal.gdc.cancer.gov/projects. The author(s) also would like to thank the GTEx, TCGA, UCSC Xena, TISCH data portal and GEO databases for the availability of the data.

## Abbreviations

AML: Acute myeloid leukemia
APCs: antigen presenting cells
AUC: Area under the curve
BM: Bone Marrow
DEGs: differently expressed genes
DFS: disease-free survival
FLT3: fms related tyrosine kinase 3
GTEx: Genotype-Tissue Expression
GEO: Gene Expression Omnibus
GEPIA: Gene Expression Profiling Interactive Analysis
GSEA: Gene set enrichment analysis
GO: Gene ontology
HLA: human leukocyte antigens
HSC: Hematopoietic stem cell
ICD: immunogenic cell death
ICPs: immune checkpoints
KEGG: Kyoto encyclopedia of genes and genomes
LASSO: Least absolute shrinkage and selection operator
MHC: Major Histocompatibility Complex
NPM1: nucleophosmin 1
OS: overall survival
ROC: Receiver operating characteristic
ssGSEA: single-sample gene set enrichment analysis
TAAs: Tumor-associated antigens
TARGET: Therapeutically Applicable Research to Generate Effective Treatments
TCGA: The Cancer Genome Atlas
TISCH: Tumor Immune Single-cell Hub
TME: Tumor microenvironment
TIMER: Tumor Immune Estimation Resource
UCSC: University of California Santa Cruz
WGCNA: weight gene co-expression network analysis.

## Notes

### Competing Interest Statement

The authors have declared no competing interest.

