## Supplementary Figures for "Identification of Tumor Antigens and Immune Subtypes of Acute Myeloid Leukemia for mRNA Vaccine Development"

### Figure S1

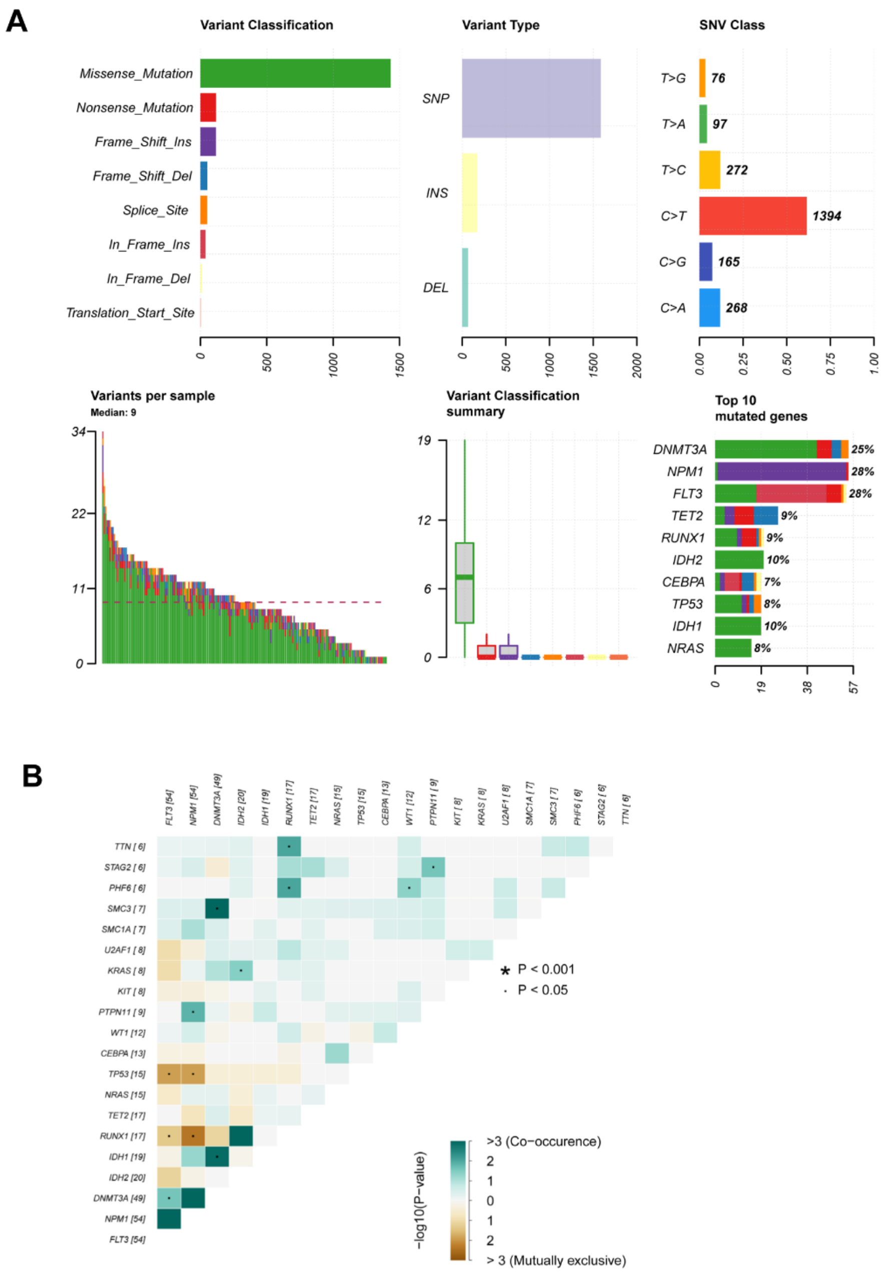


**Figure S1. Mutation analysis in TCGA-AML.** (A) Summary of mutational signature analysis in TCGA AML samples. (B) Correlation analysis of the top 20 mutated genes in TCGA AML samples.

#### Figure S2

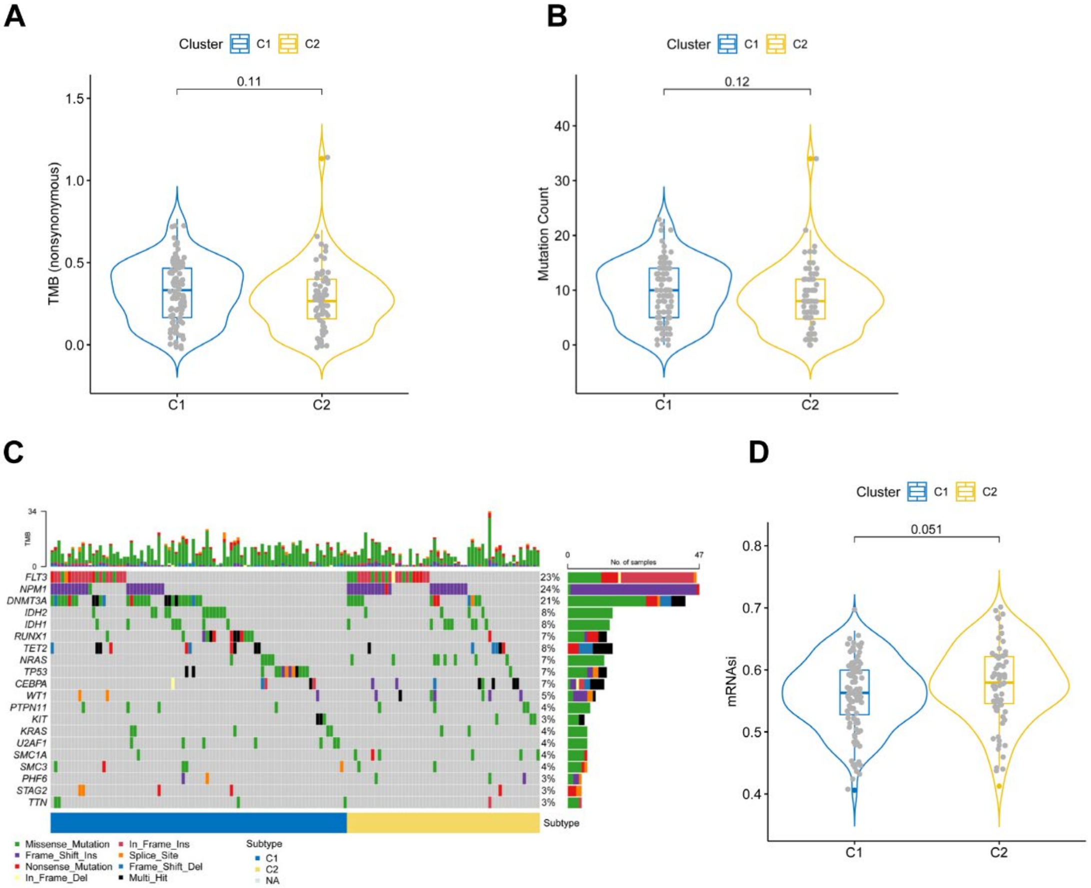


**Figure S2. Association between tumor mutation and the two immune subtypes.** (A) Comparison of tumor mutational burden (TMB) and mutation count between the two immune subtypes C1 and C2. (C) The alteration landscape of the top 20 highly mutated genes across two immune subtypes. (D) Comparison of mRNAsi in the two immune subtypes.
